# Regulation of eye movements and pupil size in natural scenes

**DOI:** 10.1101/2025.01.06.631507

**Authors:** Alexander Hahn, Aenne Brielmann, Niloufar Tabandeh, Manuel Spitschan

## Abstract

The visual diet of humans is complex in space, time, spectrum, and consequently the activation of the retinal photoreceptors. While analyses of natural scenes have yielded valuable insights, the naturalistic natural stimulus on the retina is not very well understood. In the present study, we performed eye tracking in naturalistic indoor and outdoor real-world scenes. We recorded pupil size, several saccade and fixation metrics, as well as subjective scene perception ratings derived from subjective questionnaires. For the first five seconds of eye tracking, the descriptive data analysis revealed significantly increased average saccade frequency (p =.0025), amplitude (p =.0049), peak velocity (p =.0072), as well as pupil size (p = 1.307e-09) in the indoor environments. After this initial phase, these differences vanished, except for pupil size. Using an exploratory analysis on the whole 4-minute measurement, we found that saccade and fixation metrics, along with scene ratings, showed significantly different correlations between indoor and outdoor conditions. Despite the inherent constraints of such a naturalistic study (reduced ability to exert control over the precise task and the environmental conditions), we contend that the dataset holds substantial value for field of eye movement research, as it has effectively minimized confounding factors that have been prevalent in previous eye tracking studies.

**Highlights:**

- Increased amplitude, frequency, and peak velocity of saccades in real- world indoor environments within the initial 5 seconds of eye tracking
- Different correlations of eye movement metrics and subjective scene per- ception between real-world indoor and outdoor environments
- Establishing an eye movement dataset of various saccade in fixation metrics in naturalistic real-world environments with reduced confounding factors

## 1. Introduction

The control of eye movements has been a topic of scientific discussion for decades. The vast majority of eye tracking studies were conducted in laboratory surroundings, where participants were subjected to highly artificial experimental conditions. Due to this circumstance, and the recent advent of more sophisticated eye tracking equipment, the pile of research dedicated to the characterization of eye movements under naturalistic conditions is still very small. In laboratory studies, the experimental design often includes looking at pictures of naturalistic scenes under artificial light conditions, restricted movement of the head and body, as well as large and prominent eye tracking equipment, all of which can influence the participants eye movement behavior.

Apart from the general surroundings, the tasks and instructions given to the subjects can also impact their eye movements. This was elegantly shown in pioneering eye tracking approaches by Buswell 1935 and Yarbus 1967, where differences in information (e.g., reading an article about the picture before-hand) or instruction (e.g., estimating the material circumstances of people in an image) yielded different gaze patterns. The results of more recent studies emphasize this finding by conducting eye tracking, while the participants perform specific tasks, e.g., driving, sports, walking through a supermarket, making a sandwich, navigating through a building, or working at a geological site (Kapitaniak et al. 2015, Hayhoe et al. 2012, Land 2009, Gidlö f et al. 2017, Kredel et al. 2023, Drewes et al. 2021, and Evans et al. 2012). Considering that it was not the primary goal of these studies, the number of eye movement metrics reported in their datasets is fairly limited (i.e. some studies include only the fixation number and duration while others are restricted to quantative descriptions of saccades). When evaluating these and other datasets (Henderson and Hollingworth 1999, Zwierko et al. 2023), it becomes evident that the outcome values are highly dependent on the tasks and instructions of the study protocol, which highlights the limitations of this part of the methodology in the research field of eye tracking. A naturalistic eye tracking approach is likely to minimize the influence a task might have on the acquired data set. However, the complete absence of a task presents a disadvantage as well, since the direction of high acuity foveal vision is motivated by behavioral goals (Tatler et al. 2011). Without such a goal or task, the participant is likely to generate their own intrinsic motivations and tasks, often subconscious in origin (Kay et al. 2023) and impossible to predict for the investigator. An eye-tracking approach, with the ambition of naturalistic conditions, must therefore decide carefully on this matter. In recent studies that have not imposed specific tasks on their participants, the subjects were viewing specific scenes in a more artificial context (e.g., art works in a museum) (Fontoura and Menu 2021; Stein et al. 2022). Other forms of naturalistic eye tracking involved looking at pictures of naturalistic scenes in the laboratory (Scott et al. 2020).

Moreover, as this study also focuses on pupil size measurements in the real world, one needs to consider the sluggish effect of intrinsically photosensitive retinal ganglion cells (ipRGC) (Wong et al. 2005; Berson et al. 2002) on pupil diameter. Besides the pupil size, many other non-visual functions in the central nervous system (e.g. melatonin suppression, circadian phase shifting) are affected by these cells (Spitschan 2019). In terms of their potential effect in eye movements, to the best of our knowledge no study suggested direct significant effects of ipRGCs on eye movements so far (apart from Lee and Yeh 2021), who reported reduced saccade latencies due to ipRGC stimulation). To account for the sluggish response of ipRGCs to light stimuli, a light adaptation phase would have to be implemented directly before the eye tracking measurement.

Additionally, studies in this field usually do not blind their participants in terms of their knowledge about the eye tracking approach (e.g., Drewes et al. 2021, Foulsham et al. 2011; blinding or deception are not mentioned in the eye tracking reporting guidelines by Holmqvist et al. 2022 & Dunn et al. 2023). Likely, this adds an undesired influence on the dataset, because participants can become conscious of their own gaze patterns and their potential influence on the integrity of the results. To bolster the data’s credibility, we think it is imperative to implement a procedure in which participants remain uninformed or unaware of the eye tracking, even though this is not an easy task considering they are still obliged to wear the tracking equipment.

All in all, the mentioned approaches are strongly confounded by physical restriction, task-dependency, artificial lighting conditions, general laboratory surroundings, knowledge of the subjects about the study procedure, and/or not accounting for sluggish ipRGC responses. Although certainly much knowledge has been uncovered about how the eye moves according to certain stimuli, it is crucial to consider whether these results translate to the naturalistic conditions of the real world (Foulsham et al. 2011). In the present study, we aim to contribute to this field by performing eye tracking in a set of naturalistic real-world scenes, where we compare indoor and outdoor settings in terms of various eye movement metrics and scene perception ratings. The two types of eye movement analyses we performed were [1] over the entire eye tracking duration (4 minutes) as well as [2] for the initial phase of scene viewing to examine differences in eye movements in the exploration phase of real-world scene perception. If and how eye movement metrics change within this initial phase is of particular interest. It has been shown in laboratory studies that humans extract a great wealth of scene information within the first moments of scene exploration (Anderson et al. 2022). Here, we apply a participant-blind procedure, accompanied by a substantial light adaptation phase preceding each measurement, in the absence of the aforementioned issues inherent to a particular task.

## 2. Methods

As a reference for establishing a complete and comprehensive methodology section, we drew upon the minimal reporting guidelines for eye tracking research by Holmqvist et al. 2022 and Dunn et al. 2023.

### 2.1. Participants

#### 2.1.1. Study sample

The study sample consisted of 18 participants (13 females; 24.2 mean age (range 20–29 years)) who gave their full consent for data usage after the debriefing. The mean refractive error of all participants amounted to -1.1 (dpt) (range -3,75 dpt - 0 dpt). The larger number of female participants was attributable to the larger amount of participation applications by females. Although we aspired to use a numerically equal gender distribution of participants, we do not deem data quality to be jeopardized by this. Coors et al. 2022 found that gender differences in certain eye-movement characteristics exist (blink rate, smooth pursuit gain) but can be mostly neglected in magnitude.

##### 2.1.2. Recruitment

The first instance of participant recruitment was self-selection through advertisements, which included the attachment of flyers in Tü bingen and a cir- cular mail through the newsletter of the University of Tü bingen. Individuals interested in participating in the study were asked to fill out an online questionnaire on REDCap, an online tool for data collection tool, to assess inclusion and exclusion criteria. If eligible for the study, participants were then contacted by the experimenters to agree on possible dates for the experiment to start and to discuss any further questions about the experiment. Participants were compensated at the end of the study according to their compliance with the experimental procedure: for every hour that the volunteers participated in the study procedure (measurement session + preceding screening), they received €10. Participants were remunerated, and they agreed to make their eye movement data accessible or not.

##### 2.1.3. Screening procedure

Eligible participants were selected according to the inclusion and exclusion criteria based on mental and physical health information. The age limit of the adult participants was 40 years due to the increase in optical density of the lens with age (Pokorny et al. 1987), which causes different spectral sensitivities of retinal photopigment melanopsin (Spitschan 2019). Participants were required to have normal binocular vision and color vision, as well as normal or corrected to normal visual acuity between +5 and -5 diopters. People with impairments, diseases or previous surgeries of their ocular apparatus were excluded from participation in the study, because this would inhibit correct data acquisition by the eye tracking equipment. Moreover, individuals suffering from diagnosed sleep or psychiatric disorders were excluded from participation. Sleep deprivation results in an increased saccade latency, a reduction in saccade peak velocity and smooth pursuit velocity, as well as more antisac-cade errors (Ahlstrom et al. 2013; Fransson et al. 2008; Meyhö fer et al. 2017). Moreover, it has been reported that the mental fatigue caused by sleep deprivation affects many different cognitive areas, such as cognitive speed (Dongen and Dinges 2005) and arousal (Gunzelmann et al. 2007), which in turn could affect the movement of the eyes. Additionally, sleep quality was assessed via self-report on the (PSQI) (Pittsburgh Sleep Quality Index) questionnaire. Participants that exceeded a total PSQI score of 5 were excluded from study participation.

Moreover, several psychiatric disorders have been found to alter eye movements (Katsanis et al. 1997), which is why participants with diagnosed psychiatric disorders were excluded from study participation.

The intake of any drugs and/or medications known to influence photosensitivity, movement of pupils or the ability to concentrate will be considered a criterion for exclusion. All the above-mentioned criteria for inclusion and exclusion will be assessed by self-report through an online platform developed for this purpose on REDCap. This included the standardized questionnaires (AUDIT) and PSQI, which assess alcohol consumption and sleep quality.

#### 2.2. General measurement procedure

Eligible participants were invited to the institute in teams or individually, depending on their individual time availability. During July 2023, 26 appointments (16 solo appointments and 10 team appointments) were performed over the course of 16 days. Each participant was invited to the Max-Planck Institute for Biological Cybernetics twice. A single appointment involved three measurements, either within an indoor or outdoor setting. This procedure was undertaken for a multitude of reasons. First, all conceivable scene orders were employed to mitigate the influence of sequential order on scene sequences within an individual session (3! = 6 possible scene sequences). This would not have been possible if all scenes were performed at the same appointment. Second, outdoor environments are particularly likely to decrease the sensitivity of ipRGCs to light stimuli (Wong et al. 2005) during the respective other condition, since they are usually accompanied by higher photopic illuminances. Wong et al. 2005 found that M1 ipRGCs require several hours to fully regain their sensitivity after intense light stimulation. Outdoor measurements were performed in the afternoon between 2pm and 3:30pm (CEST) (Ø*t* = 02:55pm (*±*21,2 min)). Due to time constraints, indoor measurements had to be performed in the morning between 08:30am and 10:30am (Ø*t* = 09:25am (*±*23,6 min)). We estimate that the difference in time of day has minimal effect on the task performance, because during both times the attention and alertness levels are similar during the circadian rhythm (Valdez 2019).

At the start of the first appointment, all participants went through a short screening, which included the (HRR)-test, Titmus-Fly test and Landolt-C test to exclude the presence visual malfunctions that the participants were unaware of. Additionally, prior to the first measurement, the participants performed a dim orange light adaptation phase (t = 10 minutes; *>*1 lx; 1500–1000 K), which was immediately followed by a 4-minute reference eye tracking measurement under the same dim orange light conditions. After the reference measurement and in between measurements, the participants kept wearing the eye tracker to further accustom them to the device.

Subsequently, the participants and the experimental investigators went to the first measurement location. Upon arrival, the participants faced the scene for 5 minutes prior to the start of the measurement to enable light adaptation. The adaptation phase accounted for the delayed contribution of ipRGC influence on the pupillary light reflex (McDougal and Gamlin 2010) and the saccade behavior (Lee and Yeh 2021). During this adaptation phase, the participants wore a custom-made face shield that transmitted the light to the eye but blurred their vision. This approach enabled sufficient light adaptation and simultaneously maintained a novel experience of the scene at the start of the measurement. The face shield used a diffusion material manufactured by ”Luminit” (Light Shaping Diffusor - Polycarbonate 0.010” - Angle: 40°). The shielding material transmitted 85 to 95 % of irradiated light over the entire visible spectrum. After the 5-minute adaptation phase had ended, the face shield was taken off, and the participants viewed the novel scene for 4 minutes. Before the 4-minute measurement could be initiated, however, the eye tracker had to be calibrated, which usually took 25–40 seconds. Light measurements with a Spectroradiometer were taken 10 seconds after the eye tracking measurement had started, and high-definition video capturing was performed continuously over the whole 4-minute period. After each measurement, the measurement set-up was disassembled, and the experimental conductor and the participants walked to the next measurement location.

#### 2.3. Deception approach

Before and during the data collection, participants were withheld certain information about the scope of the study. Participants were aware of the monitoring of their pupil size, but not of the tracking of their eye movements. This served the purpose of mitigating the impact of behavioral monitoring awareness on the movement of the eye, as knowledge of the very presence of an eye tracker could be influential on the subject’s eye movements. Such an impact might originate from the subject’s anticipation of the investigator’s expectations of the eye tracking results, or from what the subject feels to be socially appropriate to look at during the sessions (Risko and Kingstone 2011; Nasiopoulos et al. 2015). Unlike eye movements, which are often consciously performed, the pupil size is controlled exclusively by the autonomic nervous system and is thus not influenced by knowledge about monitoring. Correspondingly, all advertisement material, information sheets during the screening procedure, and instructions during the measurement procedure did not mention the tracking of eye movement, but only informed the participants about the monitoring of pupil size while they viewed different outdoor and indoor scenes.

After the post-experimental inquiry had been filled out, participants were debriefed about the deception immediately. The face-to-face debriefing included a list of all collected data types, as well as a detailed explanation why the deception approach was deemed necessary by the researchers. During the debriefing, the participants were asked to make their eye tracking data accessible for data analysis.

#### 2.4. Study task

In the present study, we gave the participants a naturalistic behavior task according to the definition of Kay et al. 2023. The participants were not given concrete instructions (e.g., where to look or to what to pay attention to) but were told to view the scene in a natural and unrestricted manner. Further task restrictions imposed by the experimenter would be detrimental to the aim of the study, which is to characterize natural eye movements in naturalistic real-world environments.

#### 2.5. Outdoor and indoor environments

Participants were exposed to a total of six real-world environments (ME) which consisted of three indoor environments (IME) and three outdoor environments (OME) (Fig. 1). At all experimental sites, the exact places and orientations where the participants and the measurement equipment had to be positioned were marked with tape to prevent changes between measurements. The IMEs consisted of three scenes in the Cyberneum building of the Max-Planck Institute for Biological Cybernetics in Tü bingen. In all three IMEs, all available ceiling lights were turned on during the measurements to maximize artificial lighting. While IME 1 (basement) did not include windows, IME 2 (hallway) included one window, and in IME 3 (office) several windows were present directly in the view of the participants. This gradient of naturalistic lighting in the scenes increases the diversity and, thus, the naturalistic character of the set of IMEs. If windows were included in IMEs, these were all double-glazed and kept closed and uncovered.

**Figure 1:**
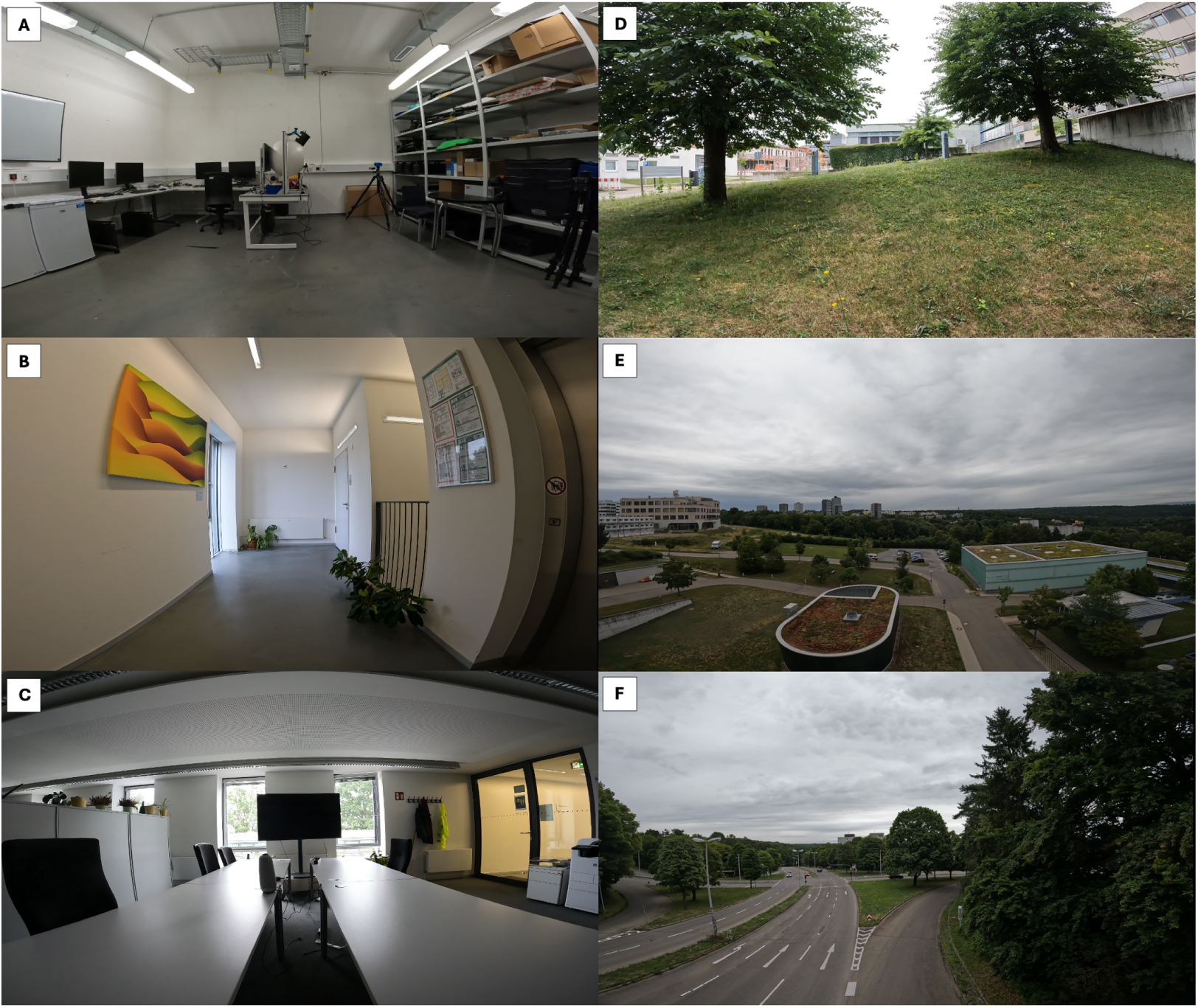
GoPro Hero 10 images (Lens mode: wide) of all measurement environments. The images only represent the part of the scene that was directly in front of the participants. Their unrestricted head movement potentially enabled them to view parts of the scene that cannot be depicted in these images.

The OMEs represented a set of exemplary naturalistic environments people in and around Tü bingen could be exposed to on a daily basis. The field of vision in all OMEs is composed largely of vegetation and artificially created objects like roads and buildings. The views towards the horizon (OME 2) and towards the building of the Max-Planck Institute (OME 1) were suspected to include only a very small number of moving objects (people, cars, bicycles, etc.) in proximity to the participants. This circumstance would likely produce a reduced number of slow-pursuit eye movements, resulting in a larger sample time for potential saccade movements and fixations. However, the goal of this study is to characterize eye movements in naturalistic environments, and since humans are exposed to a large amount of moving objects throughout their day, we deemed it appropriate to include OME 3, a view that would emulate this scenario.

#### 2.6. Measurement instruments

The spectroradiometer and the camera were mounted on a fixed holder that was positioned at the scenes with a tripod. This was done in order to guarantee the same geometry between instruments for every measurement.

##### 2.6.1. Eye tracker

Eye movement and pupil size tracking were done with the Tobii Pro Glasses 3, a video-based eye tracker providing gaze coordinate data. The build and weight of this head-mounted eye tracker are fairly similar to those of regular glasses. Thus, the influence on the natural viewing behavior is presumably reduced during the measurements in comparison to older generations of mobile eye tracking devices. The Tobii Pro Glasses 3 are pupil-centered corneal reflection (P-CR) eye trackers with eight infrared-illuminators and two cameras in each lens. Both cameras take alternating samples at a rate of 50 Hz, amounting to an effective sampling rate of 100 Hz. Eye positions were recorded for two eyes simultaneously. To perform reliable eye tracking with participants that had an optical refractive error, we used product-specific corrective lenses with optical powers ranging from +5 diopters to -5 diopters. The eye tracking device was calibrated before every measurement with the one-point calibration method provided by the manufacturer. The calibration accounted for slippage (movement of the device while wearing) and the changing light settings in between scenes. Slippage was reduced with a thin, adjustable headband attached to the frame of the glasses. The calibration procedure involved looking at a standardized calibration card that is held up to eye level at a distance of 50–100 cm from the cornea. To maximize the success of calibration and the subsequent accuracy and precision of the eye tracker, we used nose pads provided by Tobii to position the eyes as centrally in the lenses as possible. The eye tracker provides a spatial accuracy (difference between the true gaze position and the gaze position reported by the eye tracker) of 0.6°and records a field of view of 95°horizontally and 63°vertically.

##### 2.6.2. Spectroradiometer

For the purpose of this study, the spectroradiometer MSC15 from Gigahertz-Optik was used to capture spectral irradiance and illuminance. During data acquisition, the calibration certificate of the device was active, meaning the device is producing accurate results with high confidence.

#### 2.7. Questionnaires

##### 2.7.1. Post-measurement questionnaire

After each eye tracking measurement, the subjects filled out a questionnaire (see appendix). The questionnaire allowed the assessment of subjective traits of the scene (Ratings between 1 and 10), which included beauty, brightness, visual comfort and complexity. Moreover, the Karolinska Sleepiness Scale (KSS) was implemented in the questionnaire to quantify the participants.

##### 2.7.2. Post-experimental questionnaire

The effectiveness of the deception was assessed via post-experimental inquiry (see appendix) right after the last measurement had ceased. The questionnaire was answered on a tablet rather than verbally in a face-to-face interview, because this is likely to increase the honesty of the participants in such inquiries (Blackhart et al. 2012). The participants were asked about their experiences during the study, whether they knew what types of data were collected, and if they took part in cross-talk (participants divulging important experimental details with other individuals who may participate in the study). Cross-talk could have been detrimental to the study procedure if one participant had already been debriefed and the other participant had not performed all measurements yet. Before answering the post-experimental inquiries, the experimenter highlighted the scientific approach and integrity of this study, because such a statement is also suggested to increase the truthfulness of the given answers.(Blackhart et al. 2012).

#### 2.8. Data Processing and Analysis

Due to a loss of spectroradiometer data, the original number of 18 datasets was reduced to 16 for the OMEs and 17 for the IMEs. Overall, we collected 15 complete datasets of that included all eye tracking and light measurements. Furthermore, the percentage of successful samples taken by the eye tracker varied dramatically, depending on the intensity of solar irradiance, the usage of nose pads, and inter-individual differences among the participants that could not be specified in more detail. Specifically, high solar irradiance in outdoor scenes had a negative impact on eye tracking data quality. This is suggested by the fact that not one measurement had to be discarded due to insufficient eye tracking sample success rate (ETSSR) (*<*70%) from the generally darker IMEs, while in the brighter OMEs, a total of 9 measurements (out of 48) had to be discarded due to a low ETSSR. The process of discarding insufficient eye tracking data led to 14 usable datasets in OME 1, 13 usable datasets in OME 2, and 12 usable datasets in OME 3 that were used for analysis.

The eye tracking recordings were processed with the Tobii Pro Lab Software (Version 1.217), which allows direct extraction of the eye tracking data. For this, the gaze filter ”Tobii I-VT (Attention)” was used, which filters out fixations with durations *<*60 ms, since these are mostly misinterpretations of the system due to unstable pupil recognition. The data was then analyzed, using a custom code written on PYTHON.

Within the field of view that was recorded by the eye tracker, it performed measurements of the velocity, amplitude and frequency of saccades, the duration and frequency of fixations, as well as the average size of the pupil. The comparison of these metrics was performed between the conditions indoors and outdoors, as well as between all scenes. An analysis of variance (ANOVA) was performed for each metric to test for statistically significant differences between the scenes.

In the analysis of the general conditions indoors and outdoors, all the afore- mentioned metrics were tested for statistically significant differences via an independent samples t-test and the effect sizes were quantified using Cohen’s d-value. Since the ANOVA included 5 statistical tests for each ME, the typical *α*-threshold value of 0.05 for statistical significance was corrected to 0.01 according to Bonferroni to account for the accumulation of Type I errors.

Analysis of the initial 15 seconds of eye tracking included binning them into three 5-second phases. For each bin, an independent samples t-test was applied, comparing the general indoor and outdoor environment for each fixation and saccade metric.

The second topic of interest was the investigation of pairwise correlations between all analyzed metrics (eye tracking, light and post-measurement questionnaire responses) by performing a ranked correlation analysis according to Spearman. In addition, we tested whether the correlations showed statistical significance (*α* = 0.0045 (Bonferroni corrected)) by compiling the p-values for each correlation. This analysis for statistical significance was performed for the indoor condition, the outdoor condition and the combined conditions.

In the post-experimental inquiries, participants assessed their experience of the experiment. Although the loss of the of spectroradiometer data led to the exclusion of participant data in some parts of the eye tracking analysis, this analysis includes all 18 participants. This is the case, since the three affected participants also added eye tracking data to either the indoor or outdoor conditions, making their experience of the experimental sessions relevant for analysis. Additionally, we assessed whether the participants were oblivious to the deceptive approach of the study design.

Lastly, it was analyzed which parts of the scenes the participants were looking at in a naturalistic task. This was approached by defining a specific set of areas of interest (AOI) for each scene and analyzing the eye tracking data in a qualitative fashion. For that, at each scene, a snapshot with a GoPro (Lens mode: Wide) was taken that was then used to apply the automated gaze mapping function of the Tobii Pro Lab Software. This allowed for quantification of how often and for how long certain areas of the image were gazed at in total. The plotted results represent averages for all participants. The relative size of the different AOIs (table in the appendix) was calculated with a custom-written script in PYTHON that divided the number of pixels with the specific RGB color code of each AOI (opacity of AOI coloring = 100%) with the total number of pixels in the image (Fig. B.7). Thus, statements about the duration and frequency of fixations in relation to the sizes of the AOIs could be made. Testing for potential correlations according to Spearman between the relative sizes of the AOIs and the frequencies and durations of fixations was performed with the stats.spearmanr()-function from the ”scipy.stats” PYTHON package.

Ratings acquired from the post-measurement questionnaire were tested for statistically significant differences via an ANOVA (*α* = 0.01 (Bonferroni corrected)). A post-hoc analysis utilizing Tukey’s Honest Significant Difference (HSD) test was conducted to discern specific statistical significances (*α* = 0.01) between scenes, should they exist. Moreover, indoor and outdoor conditions were tested for significant differences (*α* = 0.05). For this, Welch’s t-test was applied to account for the difference in sample size between the indoor and outdoor measurements, due to the exclusion of participants from the analysis (based on data loss and ETSSR *≥* 70 %).

#### 2.9. Ethical approval

This study was reviewed and approved by the TUM Ethics Committee (approval no. 2023-231-1-S-KH).

### 3. Results

#### 3.1. Comparison of eye tracking metrics between conditions

Statistical analysis of the initial 5 seconds of eye tracking revealed signifi- cantly increased saccade metrics and pupil size in the condition ”Indoor” (IME 1-3) in comparison to ”Outdoor” (OME 1-3) (”Average Frequency of Saccades” p =.0072, d = 0.60; ”Average Amplitude of Saccades” p =.0049, d = 0.63; Average Peak Velocity of Saccades p =.0025, d = 0.68; Average Pupil Size p = 1.307e-9, d = 1.49) (Fig. 3A). None of the fixation metrics displayed similar differences between indoor and outdoor conditions. While the pupil size remained significantly larger indoors, the differences between indoor and outdoor saccades disappeared in subsequent bins.

**Figure 2:** Exemplary depiction of the set-up and a participant in IME 2 (A & B) and OME 1 (C & D). A and C show the participant during the light adaptation phase, during which he wears the face shield to allow retinal adaptation but diffuses details of the scene. In B and D, the measurement is performed. In OMEs, participants wore a hat to shield themselves and the eye tracker from direct sunlight. The blue arrows and red circles on the ground indicate standardized distances and positions..

**Figure 3:**
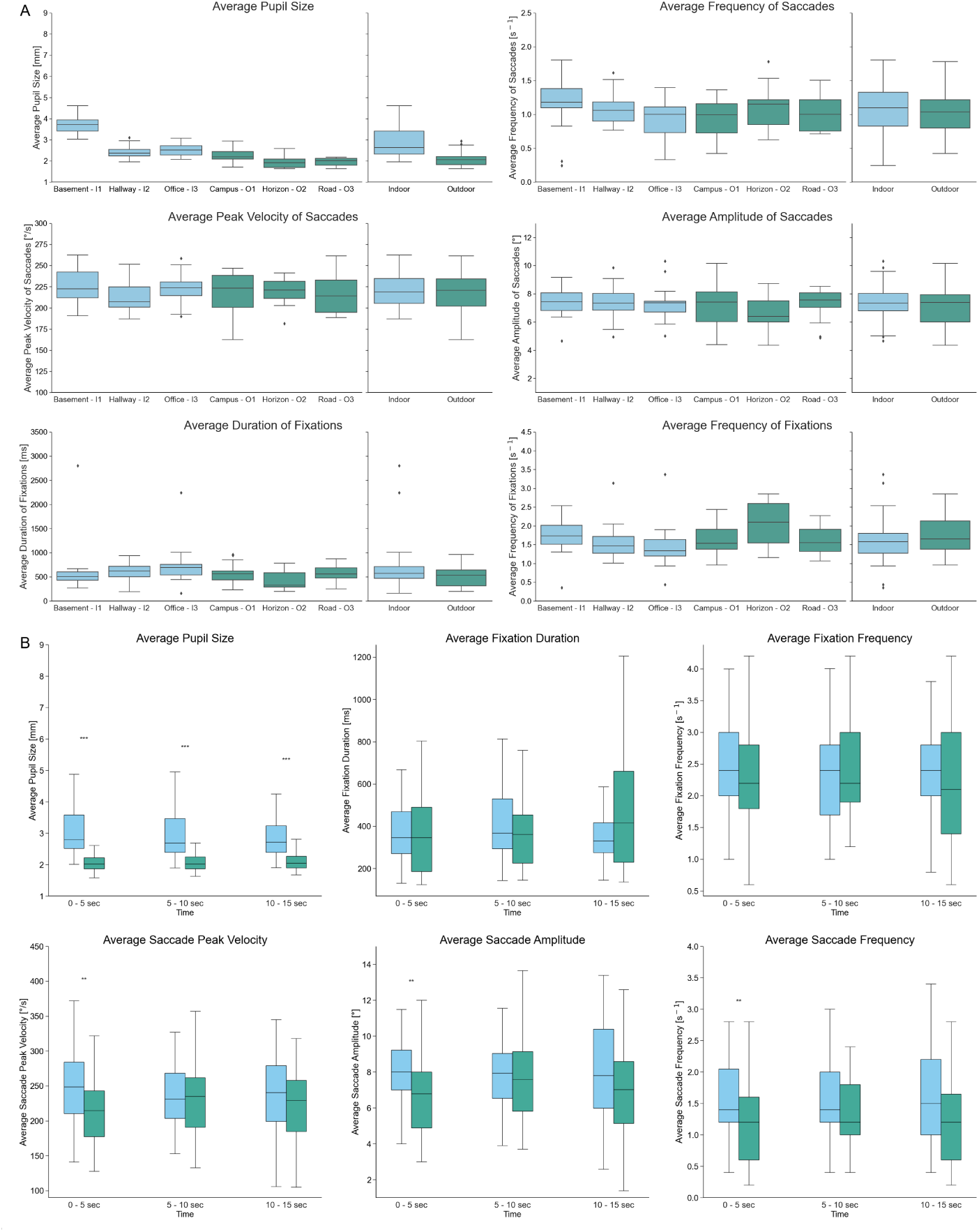
Visualization of all analyzed gaze metrics (average frequency of saccades, average pupil size, average peak velocity of saccades, average amplitude of saccades, average dura- tion of fixations, average frequency of fixations). A: Each Subplot depicts a particular metric in all six MEs and in the summarized indoor and outdoor conditions. B: Each Subplot depicts the data of the first 15 seconds of a particular metric in the summarized indoor and outdoor conditions. The time frame is fragmented into three 5-second bins..

Similarly, data analyses over the entire 4-minute measurement duration revealed no significant difference in saccade or fixation metrics between the conditions. The only exception was the ”Average Pupil Size” (Mean pupil size in = 2.88*±*0.71 mm; Mean pupil size out = 2.07*±*0.32 mm; p = 2.417e-9, d = 1.41), which was significantly increased indoors throughout the whole measurement period. None of the other metrics ”Average Peak Velocity of Saccades” (Mean APVS in = 220,41*±*19.91 °/s; Mean APVS out = 217,80*±*21.16 °/s; p =.549, d = 0.13), ”Average Amplitude of Saccades” (Mean AS in = 7.35*±*1.17 °; Mean AS out = 6.94*±*1.39 °; p =.128, d = 0.33), ”Average Frequency of Saccades” (Mean FS in = 1.07*±*0.35 s*^−^*^1^; Mean FS out = 1.03*±*0.30 s*^−^*^1^; p =.553, d = 0.13), ”Average Frequency of Fixations” (Mean FF in = 1.59*±*0.53 s*^−^*^1^; Mean FF out = 1.73*±*0.53 s*^−^*^1^; p =.206, d = 0.31) or ”Average Duration of Fixations” (Mean DF in = 653.73*±*421.11 ms; Mean DF out = 524.0*±*214.45 ms; p =.083, d = 0.37)) displayed a statistical significance difference between the conditions.

Comparisons of all six measurement environments over the entire 4-minute measurement duration (ANOVA) resulted in no statistically significant difference for ”Average Duration of Fixations” (F = 1.10; p =.37), ”Average Amplitude of Saccades” (F-value = 0.764; p =.58), ”Average Peak Velocity of Saccades” (F = 0.97; p =.44), ”Average Frequency of Saccades” (F = 1.28; p =.28) and ”Average Frequency of Fixations” (F = 2.03; p =.082) (Fig. 3B) with ”Average Pupil Size” again being the only exception (F = 63.47; p = 4.51e-27).

#### 3.2. Pairwise correlations of metrics

The pairwise correlation analysis of 11 metrics amounted to 66 correlation pairings (Fig. 4). For each pair of metrics, a separate correlation analysis of indoor, outdoor, and combined conditions was performed. The number of significant pairwise correlations amounted to 14 for the indoor condition, 10 for the outdoor condition (60% overlap with significant indoor correlations), and 17 for the combination of indoor & outdoor conditions.

**Figure 4:**
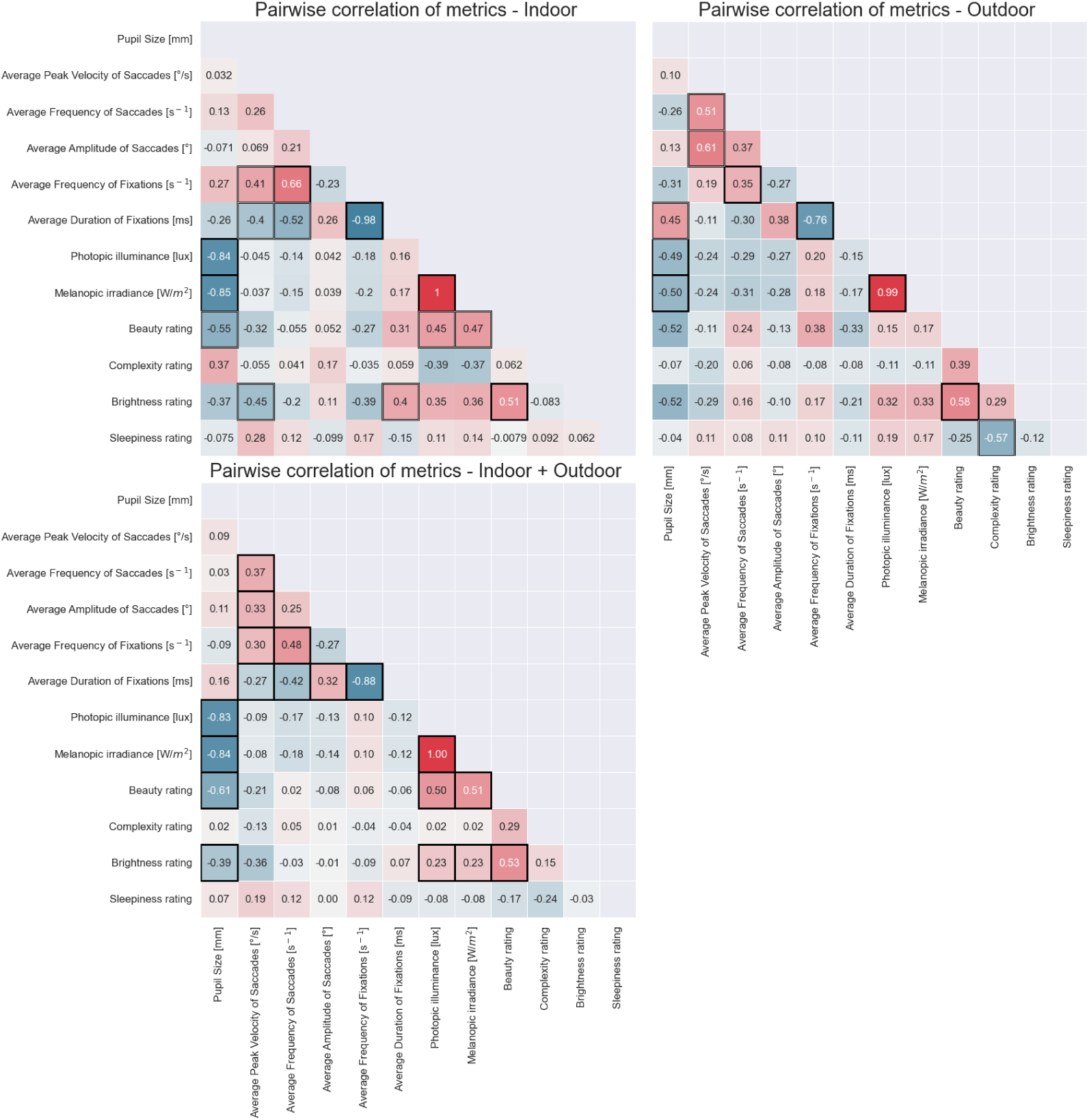
Heatmaps that visualize pairwise correlations between all analyzed metrics in three different conditions. Correlations of statistical significance are signified with a black frame around the square (Additional white framing highlights that the correlation is exclusive to either the indoor or outdoor environment)..

Apart from these overlaps, both indoor and outdoor conditions show distinct significant correlations between metrics, which will be pointed out in the following. For instance, in the indoor condition, the duration of fixations correlates negatively with the peak velocity of saccades (*ρ* = -0.40) as well as with the frequency of saccades (*ρ* = -0.52). Because duration and frequency are reciprocal metrics, the fixation frequency correlates positively with the same two metrics (*ρ* = 0.41; *ρ* = 0.66). In the outdoor condition, the average peak velocity of saccades correlates positively with the average saccade frequency (*ρ* = 0.61) and amplitude (*ρ* = 0.51). Moreover, in the outdoor condition, the average fixation duration correlates positively with the pupil size (*ρ* = 0.45). Exclusively indoors, the beauty rating of the scene correlates positively with photopic illuminance (*ρ* = 0.45) and melanopic irradiance (*ρ* = 0.47), whereas positive correlations between beauty rating and perceived scene brightness are present in both conditions (indoors: *ρ* = 0.51; outdoors: *ρ* = 0.58). Further, in the indoor condition, the perceived scene brightness correlates negatively with the average peak velocity of saccades (*ρ* = -0.45) and positively with the average fixation duration (*ρ* = 0.40).

#### 3.3. Post-measurement questionnaire

The post-measurement questionnaire was filled out by the participants im- mediately after every measurement and included several questions concern- ing their subjective experience of the measurement environment by the partic- ipants. The responses are visualized in Figure 5.

**Figure 5:**
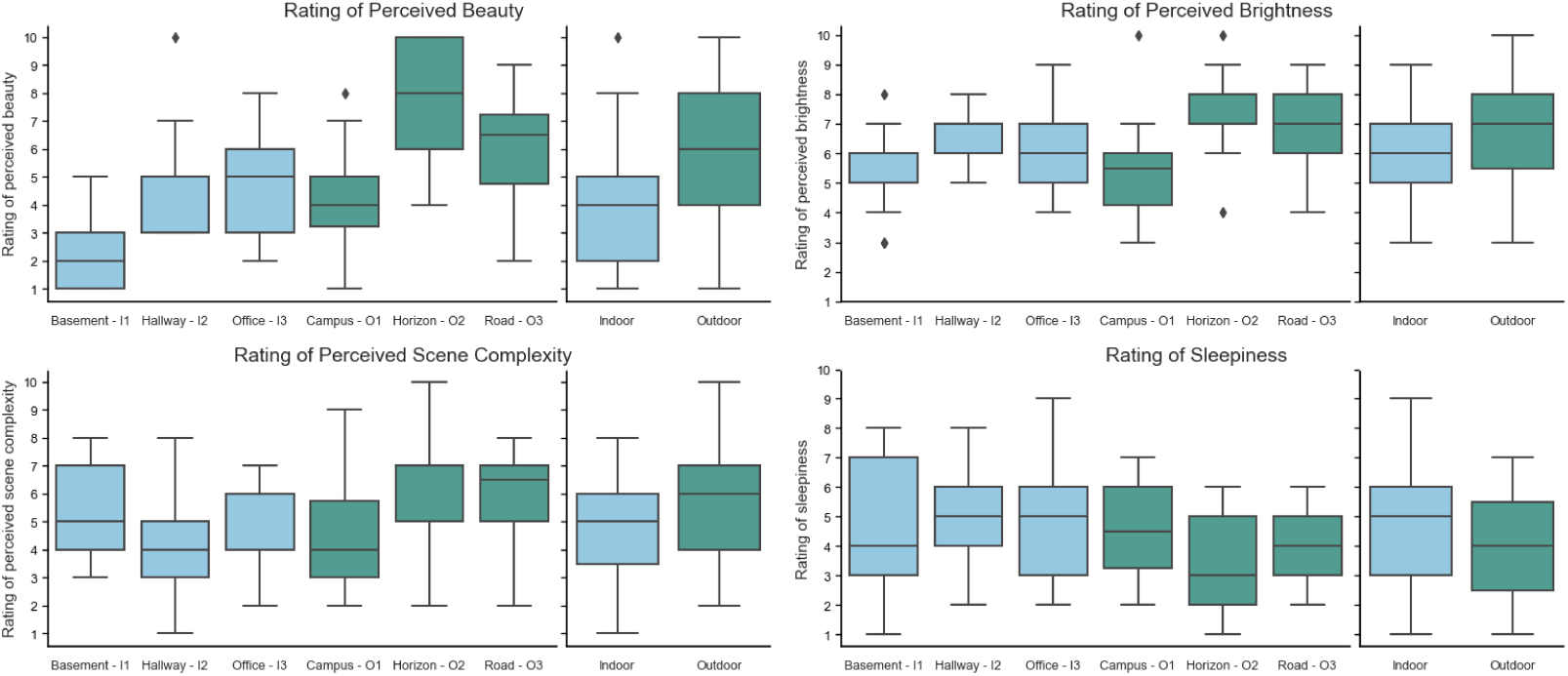
Ratings acquired from the post-measurement questionnaire regarding the ”Per- ceived Beauty”, ”Perceived Brightness”, ”Perceived Visual Comfort”, ”Perceived Complexity” and ”Sleepiness” at every measurement environment..

The results of the ANOVAs indicate significant differences in ratings between some scenes in the categories ”Beauty” (F = 13.21; p: 1.71e-9), ”Brightness” (F = 4.084; p =.0023), and ”Complexity” (F = 3.82; p =.0036). The perceived beauty was rated significantly different between indoor and outdoor scenes (Welch’s t-test: p = 5.12e-5; d = 0.93). Tukey’s HSD revealed that the highest rating of beauty was attributed to OME 2 (p-adj *<*.05); however, the difference between OME 2 and OME 3 is not statistically significant (p-adj =.128). Moreover, IME 1 was significantly less beautiful than any other scenes (Tukey’s: p-adj *<*.05).

Although the photopic illuminance and melanopic irradiance measured during the experiments varied largely between the measurement environments, the subjective differences in brightness are subtle. The only significant differences in brightness ratings arose from the comparison of IME 1 and OME 2 (Tukey’s: p-adj =.0073). In summary, the perceived brightness of IMEs and OMEs was not significantly different. Despite the significant differences in the rating of beauty, the measurement environments did not exhibit significant differences in visual comfort.

The perceived complexity of the indoor and outdoor environments was not significantly different (p =.078). The only significant difference of the complexity of two scenes originated from the comparison of IME 2 and OME 2 (Tukey’s: p-adj =.0070).

The comparison of sleepiness information in the indoor and outdoor conditions did not amount to a statistical difference (Welch’s t-test; p =.0609). Scene-specific comparisons of sleepiness (Tukey’s HSD) did not reveal any significant differences either.

#### 3.4. Light metrics

The intensity of light varied largely between the scenes and especially between the indoor and outdoor conditions (Fig. A.6). Measurement environments of the outdoor condition are generally more brightly illuminated with higher photopic illuminance (mean photopic illuminance = 8883 lux (range: 1395 lux to 56290 lux)) and a higher melanopic irradiance (mean melanopic irradiance = 8812 W/sqm (range: 1297 W/sqm to 49360 W/sqm)) in comparison to the indoor condition (mean photopic illuminance = 520 lux (range: 120 lux to 1149 lux; mean melanopic irradiance = 388 W/sqm (range: 65 W/sqm to 894 W/sqm))).

#### 3.5. Eye tracking quality in different conditions

Here, we investigated the relationship between ETSSR and the present levels of photopic illuminance during the measurements (Fig. A.6). For this analysis, only OME were considered since illuminance and ETSSR varied much more in the outdoor condition (in the indoor condition, every eye tracking measurement amounted to *>*85 %ETSSR). Outdoor measurements conducted at photopic illuminances below 10000 lux (ØETSSR = 90.54%) were significantly more successful than measurements during which the equipment was subjected to photopic illuminance above 10000 lux (ØETSSR = 77.86%) (two independent samples t-test with *α* =.05; p =.023; d = 0.68).

#### 3.6. Post-experimental inquiry

A total of five participants stated they suspected eye movement recordings during the course of the experiment. Furthermore, reports were given by five participants concerning physical discomfort and one participant about psychological discomfort during the measurements. The physical discomfort was always attributed to pain on the nose and ears caused by the pressure of the glasses, while the psychological discomfort was based on people being in the same room as the participant during the measurement. Participants who have stated that the task felt unnatural (n = 2) specified that either the four-minute measurement procedure of sitting and viewing felt unusual or wearing glasses was foreign to them. Moreover, participants were asked whether the glasses affected their viewing experience. If not answered with ”Not at all” (n = 12), they were requested to describe the effect. Most commonly, the given reason for an altered viewing experience was the weight and physical feeling of the glasses on the nose and ears (n = 3), as well as the fear of impairing the success of the measurement by larger head movements (n = 2). The post-experimental inquiry responses of all 18 participants are presented in table A.1.

### 4. Discussion

In this study, we performed eye tracking under naturalistic conditions to find potential differences between indoor and outdoor conditions. The descriptive data analysis of the recorded eye movement metrics did not yield major differences between indoor and outdoor conditions. An additional exploratory data analysis revealed several different pairwise metric correlations of significance between indoor and outdoor conditions.

#### 4.1. Comparison of eye tracking metrics between conditions

Comparison of eye movements between indoor and outdoor conditions revealed differences in saccade metrics exclusively for the first five seconds of eye tracking. This finding suggests that the initial exploration phase of the scenes differs between indoor and outdoor. The statistical analysis of gaze metrics over the entire measurement duration did not find significant differences between the indoor and outdoor conditions, as well as between the individual scenes. It is likely that the attention of initial scene exploration rapidly decreases and does not show effects in the subsequent bins.

Previous research has empirically demonstrated that tasks wield significant influence on eye movement. The aim of the study, creating conditions for unbiased eye movement behavior, could be achieved solely with a very naturalistic task and careful instructions. As expected, the metrics that we have chosen to analyze in this study deviated from the results of different studies, since every study imposes different instructions and tasks on its participants. The utilization of a naturalistic task (*”View the scene that is in front of you and feel free to look wherever you want”*) gives this dataset the advantage of reduced susceptibility to confounding by specific task instructions. While data collection with loosely defined task instructions may present additional challenges (e.g., drift of attention; emergence of intrinsic tasks or motivations), we assert that this dataset holds substantial value for the domain of eye movement research.

#### 4.2. Pairwise correlations of metrics

This exploratory correlation analysis highlights several pairwise correlations that were either exclusive to indoor or outdoor condition, or universally present in both conditions. Negative correlations between reciprocal frequency- and duration metrics will not be discussed in further detail, as their origins are evident.

Several strikingly different gaze-metric correlations between indoor and outdoor conditions were revealed by this analysis. For instance, in the outdoor condition, the ”Average Peak Velocity of Saccades” correlated positively with the other saccade metrics. In the indoor condition, this was not present, but instead the ”Average Peak Velocity of Saccades” correlated positively with the ”Average Frequency of Fixations” and negatively with the ”Average Duration of Fixations”. Another interesting finding is the rather strong positive correlation between perceived scene beauty and perceived scene brightness in both conditions implicating that the brightness of a scene is a major driver of perceived beauty. Several previous studies already showed that brightness/lightness correlates with the preference of colors (Palmer et al. 2012), but to the best of our knowledge, the present study is the first to suggest this in real-world environments.

Moreover, in all conditions, the size of the pupil shows strong negative correlations with the intensity of light present during the measurement, which was to be expected due to the pupillary light reflex.

#### 4.3. Eye tracking quality in different conditions

Generally, in an indoor environment, eye tracking was more successful than outdoors. The reasons underlying this were lower intensities of light and more stable lighting circumstances during the measurements. In the outdoor environment, eye tracking was significantly more successful at lower light levels (*<*10000 lux). Although the spectroradiometer did not measure infrared light, we assume much higher irradiances of infrared light in outdoor conditions due to the composition of sunlight. This was likely an important influence on the tracking ability of our video-based eye tracker, which illuminates the pupil with infrared light at approximately 850 nm wavelength. Furthermore, some participants reported that they squinted their eyes when subjected to high illuminance in the outdoor condition. This further reduced the ability of the pupil cameras to track the pupil.

#### 4.4. Post-measurement questionnaire

In the present study, outdoor environments were perceived as significantly more beautiful than indoor environments. This was to be expected as previous studies found that people generally believe beauty to lie more in nature than in artificial objects (Brielmann et al. 2021) and that ratings of beauty/attractiveness are increased when spatial structures of images resemble structures found in natural scenes (Palmer et al. 2012). Especially OME 2, which offered a wide-ranging view with heterogeneous content, was perceived as attractive. Contrary to that, the windowless room in IME 1 was considered to be significantly more unattractive. Interestingly, the rating of brightness did not significantly differ between indoor- and outdoor conditions, even though the measured light metrics varied drastically. The likely reason for that is the high surface reflectance of the predominantly white indoor surfaces. The visual system factors out the physical properties of the light (illumination), and the awareness shifts towards the nature of the observed objects, rather than its physical properties (proximal mode of perception).

Surprisingly, participants did not feel a significant difference in sleepiness between the conditions indoors and outdoors. The enhancing cognitive effect of bright light (Vandewalle et al. 2006, Shishegar and Boubekri 2016) and the increase in alertness by blue-enriched light (Chellappa et al. 2011) led us to the expectation of reduced sleepiness ratings in outdoor conditions.

We excluded the results from question three (*perceived visual comfort*) because three participants mentioned in the later stages of the experiment that they misunderstood the rating scale of the question. We cannot confidently estimate how many participants have had the same issue but did not point it out to the investigators.

#### 4.5. Post-experimental inquiry

The responses to this questionnaire serve to estimate the effectiveness of the deception approach and how participants were possibly distracted during the measurements. The responses showed that five out of eighteen people suspected eye tracking in this study, even though it was not mentioned to them. This could either be due to previous experiences with eye tracking studies, familiarity with the Tobii device, general suspicion of the acquired data sets, or multiple of the named reasons.

Out of 18 participants, one responded that they felt distracted by other people being in the same room as them during the measurements. This situation occurred regularly in IME 2, especially during the weekdays, suggesting that more people could have felt distracted, although they have not pointed it out in the inquiry. The subjection of other people could be considered as a significant issue in IME 2. Future studies exploring similar topics are advised to choose measurement environments where participants are not subjected to other people in the same room. A similar issue of unequal conditions due to the presence of others arose from the fluctuating number of participants that were present during the appointments. Based on their availability, participants performed the measurements either in teams of two or alone. This might have had an influence on the viewing behavior, which is little understood until now (Holmqvist et al. 2022.

### 5. Conclusion and Outlook

With this study, we set out to answer whether indoor and outdoor environments have an influence on the eye movement behavior of participants under naturalistic conditions. In summary, our findings implicate a distinctly different saccade behavior in the initial scene exploration phase, which quickly fades, while fixations display no such characteristics. Considering the whole measurement period, the two conditions significantly differ only in terms of metric correlations and pupil size. Pupil size differs vastly, which was to be expected due to the large conditional differences in measured light characteristics.

While the study aimed to minimize the confounding factors of previous eye tracking studies as much as possible, the procedure was associated with some limitations. The naturalistic character of the procedure was not able to exert task control over the participant and subjected them to conditions that were relatively uncontrolled in comparison to laboratory studies. Although this variance is part of the condition’s naturalistic character, it makes it more complicated to trace the measured results back to their physiological origin. Additionally, the chosen set of scenes was likely not broad and diverse enough to make a universally valid statement about indoor vs. outdoor conditions. To minimize this issue, we advise future studies with similar research goals to compile a larger and more diverse set of indoor and outdoor scenes, which could then potentially be assigned to categorize (e.g., three scenes for each category ”basement”, ”office”, ”living area” in the overall condition ”indoor”).

How the results of the participants-blind procedure compare to studies with emphasized knowledge of the participants (explicitly informing participants of the use of eye tracking in the study) remains unclear. The potential of too much knowledge to exert a substantial confounding effect, might be explored in future studies. Until then, we deem it sensible to adapt our deception approach for similar studies to reduce the likelihood of confounds.

To conclude, the mentioned differences between the two conditions suggest slightly different viewing behaviors in indoor and outdoor environments. In terms of the dataset itself, we assert its substantial value to the field of eye movement research, as eye movement metrics yield significantly divergent outcomes across different tasks. We hope that future studies will try to duplicate our results with a similar naturalistic approach and a different set of scenes. Given that this study applied task instructions that were specifically designed to not inhibit the naturalistic viewing conditions, we propose that the obtained results may serve as benchmark values for future studies conducting naturalistic eye tracking investigations.

## Acknowledgement

We wish to to thank Max Dobberkau, Elisa Schrö pel and Marcelo Stegmann who helped with piloting measurements, Carolina Guidolin for her helpful comments on this article and and Rafael Lazar for his valuable advice about eye tracking procedures.

### Appendix A. Additional plots, tables, questionnaires

### Appendix B. Analysis of AOI in naturalistic scenes

#### Appendix B.1. Methods Appendix B.1.1. Data Analysis

It was analyzed which parts of the scenes the participants were looking at in a naturalistic task. This was approached by defining a specific set of areas of interest (AOI) for each scene and analyzing the eye tracking data in a qualitative fashion. For that, at each scene, a snapshot with a GoPro (Lens mode: Wide) was taken that was then used to apply the automated gaze mapping function of the Tobii Pro Lab Software. This allowed for quantification of how often and for how long certain areas of the image were gazed at in total. The plotted results represent averages for all participants. The relative size of the different AOIs (B.4 was calculated with a custom-written script in Python that divided the number of pixels with the specific RGB color-code of each AOI (opacity of AOI coloring = 100%) with the total number of pixels in the image (Fig. B.7). Thus, statements about the duration and frequency of fixations in relation to the sizes of the AOIs could be made. Testing for potential correlations according to Spearman between the relative sizes of the AOIs and the frequencies and durations of fixations was performed with the stats.spearmanr function from the ”scipy.stats” Python package.

#### Appendix B.2. Results

In each ME, sets of AOIs were defined to analyze the average duration of fixations (Fig. B.8) and the frequency of fixations (Fig. **??**) in each AOI. Moreover, potential correlations between AOI size and the aforementioned metrics were investigated The only statistically significant positive correlations between average fixation frequency and AOI size were found in IME3 (*ρ* = 0.534; p =.049) and OME1 (*ρ* = 0.649; p =.0425). For all other MEs the statistical analysis did not reveal any correlation between this metric and the size of the specific area of interest. Furthermore, in neither the indoor nor outdoor ME, correlations between the average duration of fixations and AOI size were detected.

#### Appendix B.3. Discussion

Correlations between AOI size and fixations frequency / fixation duration do not indicate differences between indoor and outdoor conditions. Both have produced one significant positive correlation between AOI size and fixation frequency and none between AOI size and fixation duration. Moreover, the results suggest that AOI size is not the main factor that drives the duration of fixations, since no significant correlation with AOI size could be reported. On the other hand, AOI size might be a more important factor for an increased frequency of fixations, since two significant positive correlations in measurement environments were observed. In the absence of a prescribed directive to observe the scene with a specific task or objective, participants are likely to allow their gaze to freely traverse the scene. In this case, it might serve as an explanation for the correlation between AOI size and fixation frequency, that the participant’s unguided gaze would result in a more random gaze distribution, leading to more fixations in larger AOIs. Exceptions here would be AOIs that are visually uncomfortable (e.g., ”sky” being too bright), which are in return fixated less often. The most important factor for increased duration of a fixation is likely the complexity of an AOI (e.g., moving objects, vehicles, emission of sound). Some might argue that the frequency and duration of fixations are negatively correlated metrics. This is true if the scene is not fragmented into multiple AOIs but considered as a whole (Fig. 4). In this case, however, an AOI that is fixated for several seconds on average, can be visited very frequently as well (examples of this would be ”Windows” in IME 3 or ”Trees” in OME 1).

Although the applied method of AOI analysis in naturalistic and physically unrestricted conditions gives a sufficient proxy for which objects and structures were looked at more often than others, it also comes with a significant drawback. Fixations of AOIs could only be captured if the fixations occurred within the snapshot. Gaze patterns that were performed outside the snapshots’ framework were therefore lost and could not be included in the analysis.

---

**Figure A.6:**
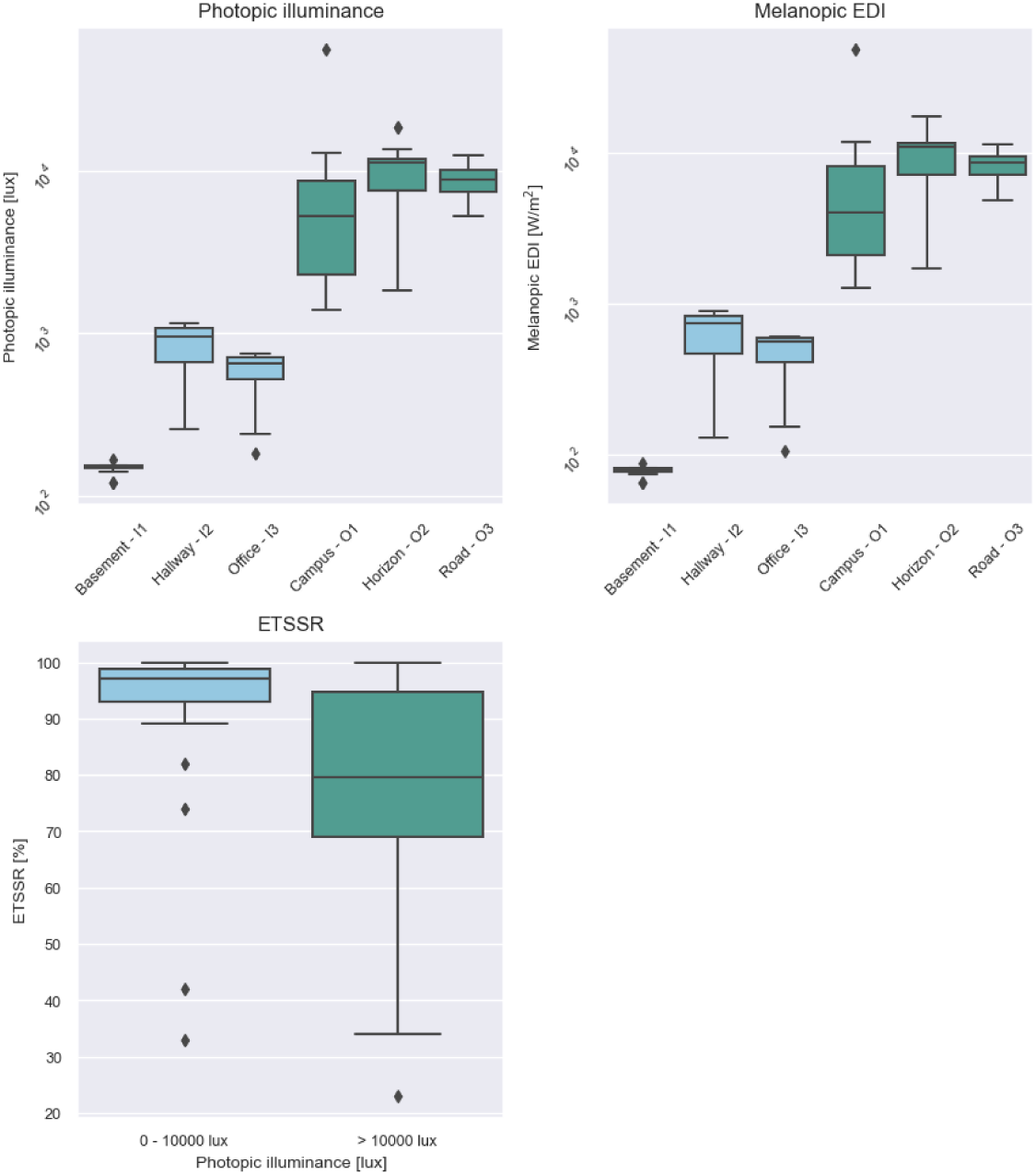
A: Photopic illuminance and melanopic EDI in the six measurement environments. B: Percentages of eye tracking sampling success rate (ETSSR) in relation to outdoor illumi- nance levels above and below/equal 10000 lux..

**Figure B.7:**
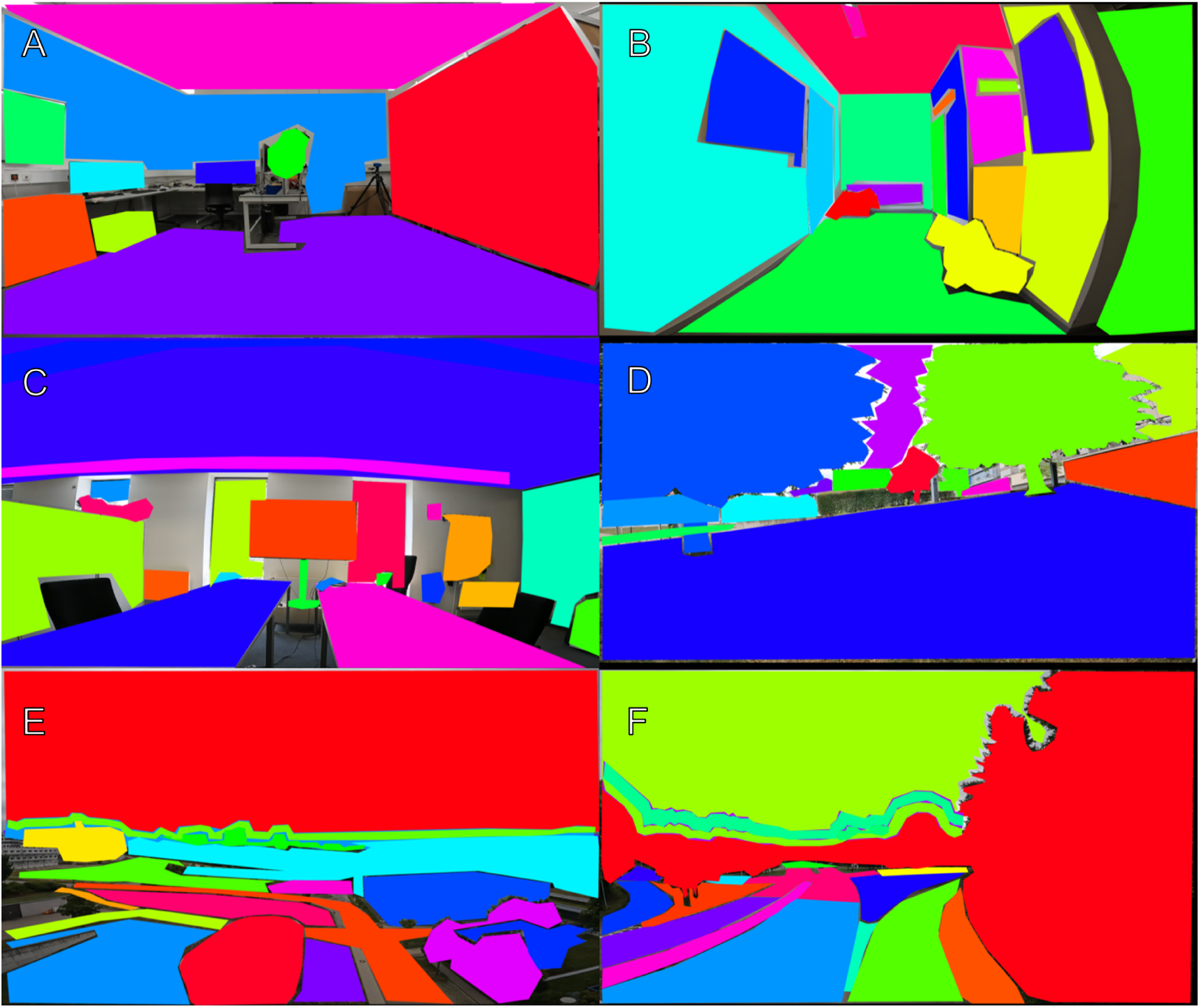
Labeling of areas of interest in all measurement environments. IMEs are displayed in image A - C. OMEs are displayed in image D - F.

**Figure B.8:**
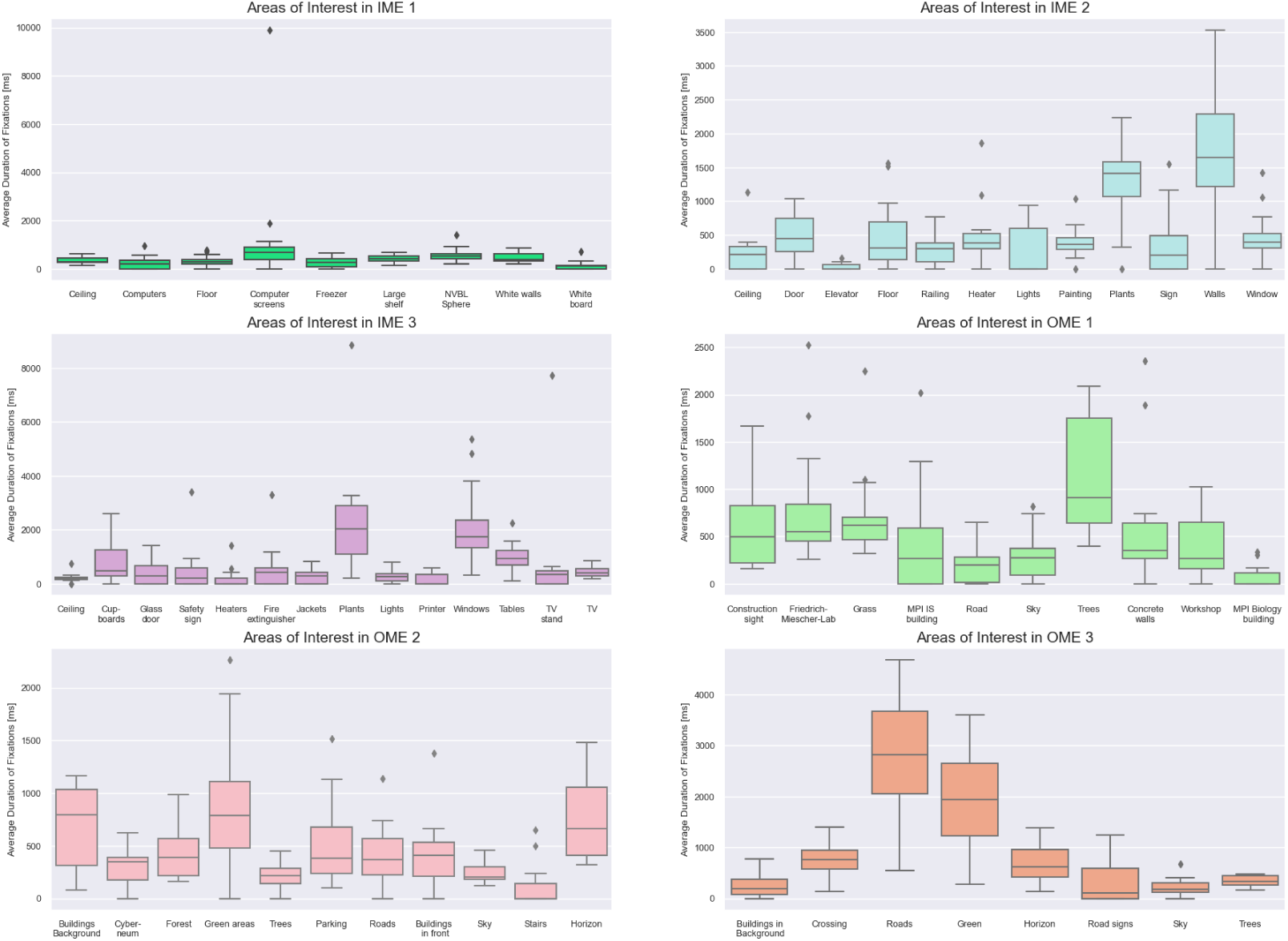
Average duration of fixation in AOIs. Each subplot represents one measurement environment. The AOIs between measurement environments differ in quantity, identity and size. The corresponding sizes of the AOIs are available in the appendix (Table B.4).

**Figure B.9:**
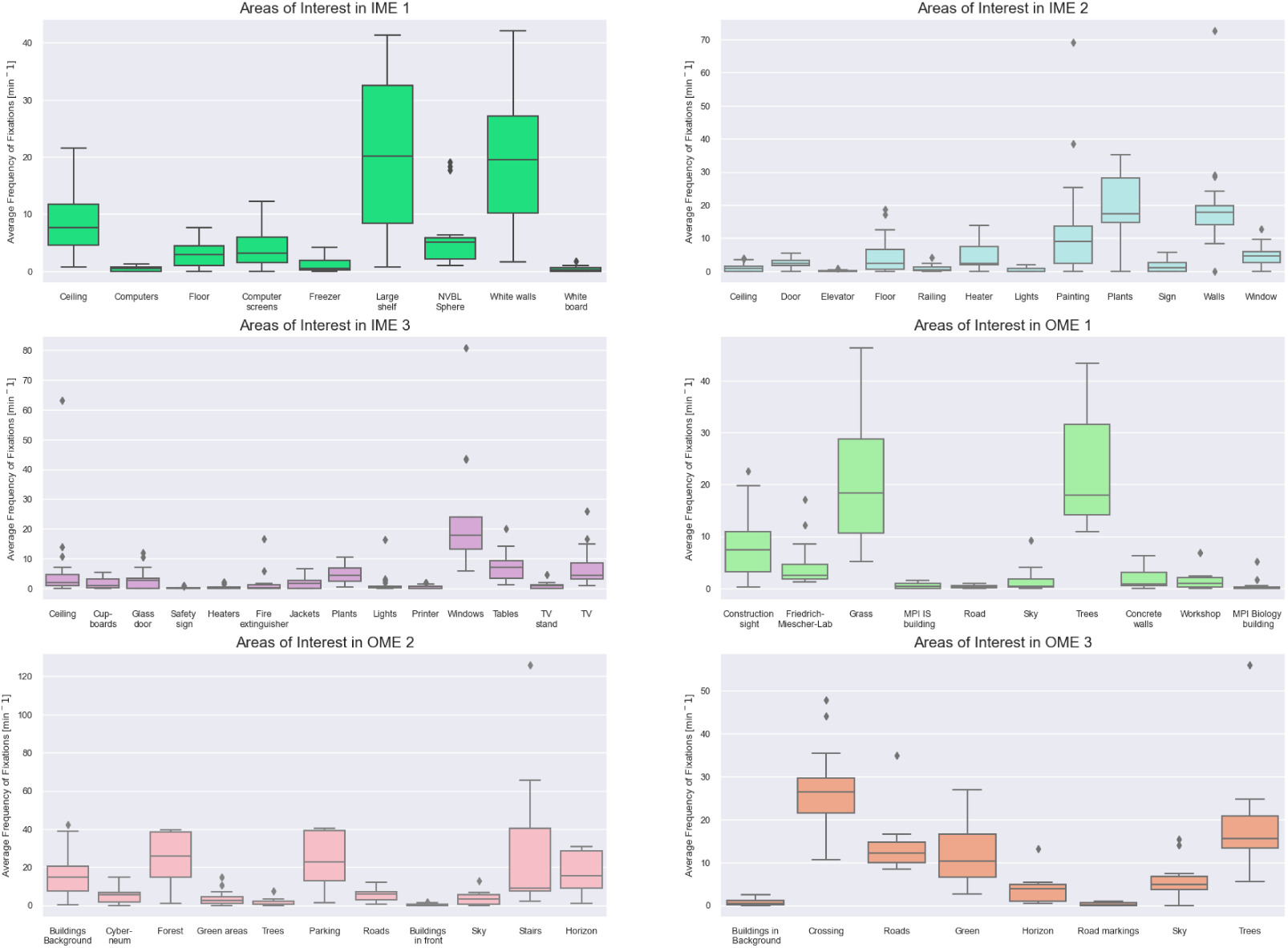
Average frequency of fixation in AOIs. Each subplot represents one measurement environment. The AOIs between measurement environments differ in number, identity and size. The corresponding sizes of the AOIs are available in the appendix (Table B.4).

**Table A.1:** Answers in the post-experimental inquiry used to assess the experiences of the participants during the experiment. Q1: Did you experience physical and/or psychological discomfort during either the first or second appointment?; Q2: To what extent do you believe that wearing the glasses modified how you looked at the indoor scenes?; Q3: How likely would you be exposed to indoor scenes similar to the ones you were exposed to in your daily life?; Q4: How likely would you be exposed to outdoor scenes similar to the ones you were exposed to in your daily life?; Q5: How natural was the task to you?; Q6: Were the instructions by the study personnel clear?; Q7: Do you know what types of data were collected throughout the study?; Q8: Are you aware of anyone you know, who participated in the study (besides your team partner)?; Q9: Did you and the other person share information about the purpose of the study?.

| ID | Q1 | Q2 | Q3 | Q4 | Q5 | Q6 | Q7 | Q8 | Q9 |
| --- | --- | --- | --- | --- | --- | --- | --- | --- | --- |
| 1 | No | Not at all | Likely | Neutral | Unnatural | Yes | Yes | No | No |
| 2 | Yes | Little | Neutral | Likely | Natural | Yes | Partly yes | No | No |
| 3 | No | Little | Neutral | Likely | Natural | Yes | Yes | No | No |
| 4 | No | Little | Likely | Very likely | Neutral | Yes | Partly yes | No | No |
| 5 | Yes | Little | Neutral | Very likely | Natural | Yes | Partly yes | Yes | No |
| 6 | No | Not at all | Neutral | Very likely | Natural | Yes | Partly yes | No | No |
| 7 | Yes | Not at all | Likely | Likely | Very natural | Yes | Partly yes | No | No |
| 8 | No | Not at all | Neutral | Very likely | Unnatural | Yes | Partly yes | No | No |
| 9 | No | Not at all | Very likely | Very likely | Natural | Yes | Yes | No | No |
| 10 | Yes | Not at all | Likely | Likely | Natural | Yes | Yes | No | No |
| 11 | Yes | A bit | Likely | Very likely | Natural | Yes | Yes | Yes | No |
| 12 | No | Not at all | Very likely | Likely | Natural | Yes | Partly yes | No | No |
| 13 | No | A bit | Likely | Likely | Natural | Yes | Partly yes | No | No |
| 14 | No | Not at all | Likely | Very likely | Natural | Yes | Yes | No | No |
| 15 | No | Not at all | Likely | Neutral | Natural | Yes | No | No | No |
| 16 | No | Little | Likely | Very likely | Neutral | Yes | Partly yes | No | No |
| 17 | Yes | A bit | Unlikely | Neutral | Neutral | Yes | Partly yes | No | No |
| 18 | No | Little | Unlikely | Likely | Natural | Yes | Partly yes | No | No |

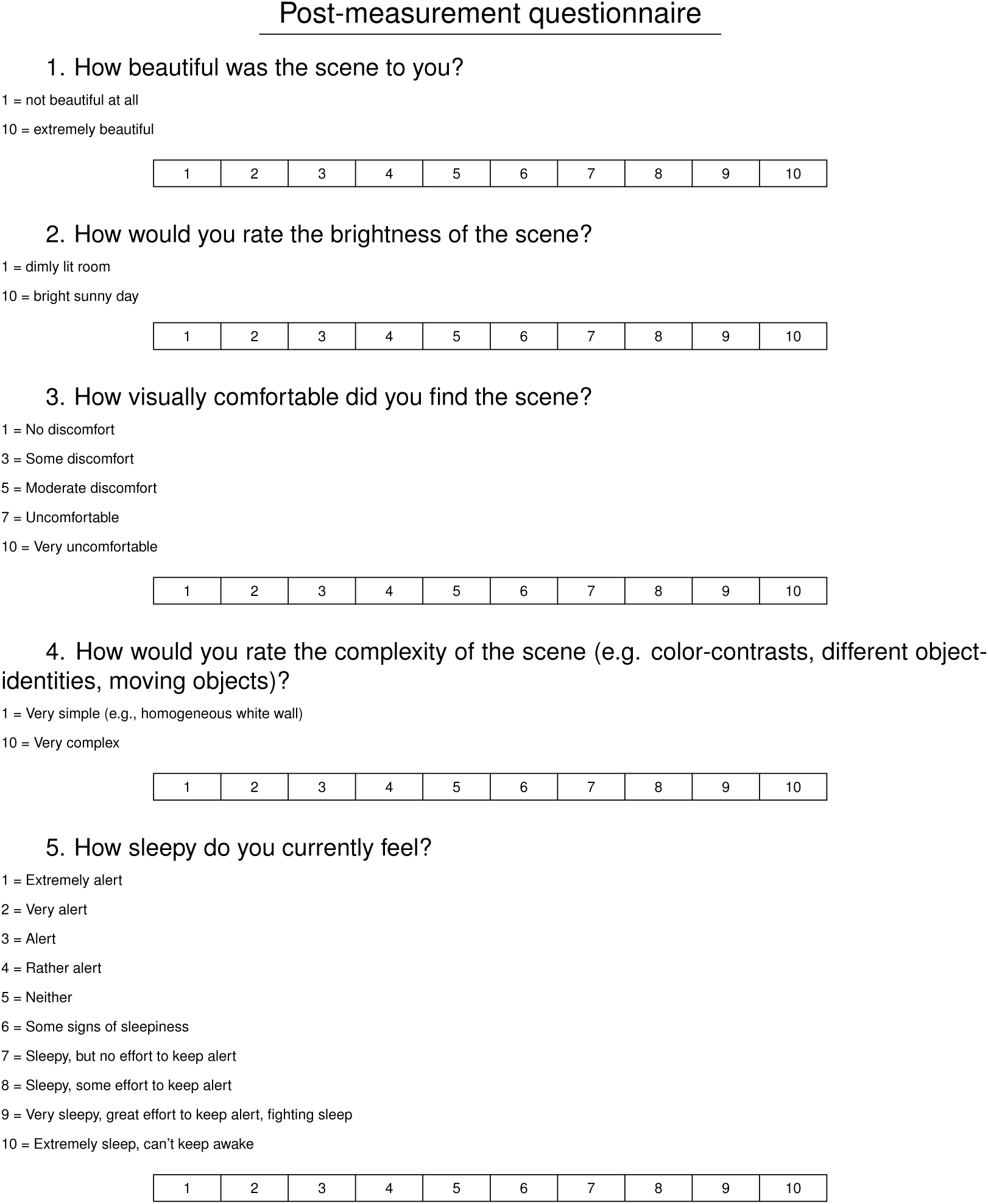

|  |  |  |  |  |  |  |  |  |  |
| --- | --- | --- | --- | --- | --- | --- | --- | --- | --- |
| 1 | 2 | 3 | 4 | 5 | 6 | 7 | 8 | 9 | 10 |
| 1 | 2 | 3 | 4 | 5 | 6 | 7 | 8 | 9 | 10 |
| 1 | 2 | 3 | 4 | 5 | 6 | 7 | 8 | 9 | 10 |
| 1 | 2 | 3 | 4 | 5 | 6 | 7 | 8 | 9 | 10 |
| 1 | 2 | 3 | 4 | 5 | 6 | 7 | 8 | 9 | 10 |

**Table A.2:** Metadata for every appointment that was conducted indoors. IME3 p.p. stands for
people present at the scene.

| Appointment | Number of participants | Date | Time | Temperature [°C] | Notes |
| --- | --- | --- | --- | --- | --- |
| 1 | One | 14.07.23 | 08:30 | n.a. | IME 3 03:45: Person walked into the field of view; IME3 p.p. |
| 2 | Two | 15.07.23 | 09:30 | n.a. | IME 2: Measurement had to be stopped and repeated after 45sec due to loss of connection |
| 3 | Two | 17.07.23 | 09:00 | n.a. | IME 3 people present (p.p.) |
| 4 | One | 18.07.23 | 09:30 | n.a. | IME 3 p.p. |
| 5 | Two | 19.07.23 | 10:30 | n.a. | OME 2&3: Calibration issues with both participants; IME 3 p.p. |
| 6 | One | 20.07.23 | 09:30 | n.a. | IME 3 p.p. |
| 7 | Two | 21.07.23 | 09:30 | n.a. | IME 3 p.p. |
| 8 | Two | 22.07.23 | 09:00 | n.a. |  |
| 9 | One | 24.07.23 | 09:30 | n.a. |  |
| 10 | One | 25.07.23 | 09:30 | n.a. |  |
| 11 | One | 26.07.23 | 09:30 | n.a. |  |
| 12 | One | 28.07.23 | 09:00 | n.a. |  |
| 13 | One | 29.07.23 | 09:30 | n.a. |  |

**Table A.3:** Metadata for every appointment that was conducted outdoors. The temperature data for Tübingen was retrospectively added to the data set via the website Meterostat 2023.

| Appointment | Number of participants | Date | Time | Temperature [°C] | Notes |
| --- | --- | --- | --- | --- | --- |
| 1 | One | 14.07.23 | 15:00 | min:13.9<br>max:28.0<br>avg:21.2 |  |
| 2 | Two | 15.07.23 | 15:00 | min:18.0<br>max:33.3<br>avg:25.5 | OME 2 02:10: Protective hat blown off by wind |
| 3 | One | 17.07.23 | 15:30 | min:15.3<br>max:25.2<br>avg:20.4 |  |
| 4 | One | 18.07.23 | 15:00 | min:13.3<br>max:25.3<br>avg:20.0 |  |
| 5 | Two | 19.07.23 | 15:00 | min:17.9<br>max:26.5<br>avg:21.9 | OME 2&3: Calibration issues with both participants |
| 6 | Two | 20.07.23 | 15:00 | min:15.9<br>max:24.1<br>avg:19.9 | OME 2: Gathering of 5-7 people ca. 10 meter behind us |
| 7 | One | 21.07.23 | 14:00 | min: 13.8<br>max:23.9 avg: 18.8 |  |
| 8 | Two | 22.07.23 | 15:00 | min:11.5<br>max:23.6<br>avg:18.3 |  |
| 9 | Two | 23.07.23 | 15:00 | min:15.6<br>max:26.5<br>avg:21.0 | OME 2&1: Calibration issues; 01:55: participant was bothered by flying insect |
| 10 | One | 27.07.23 | 15:00 | min:11.4<br>max:21.4<br>avg:16.4 | OME 1 loud noise from construction sight |
| 11 | One | 28.07.23 | 16:00 | min:14.8<br>max:23.0<br>avg:17.9 |  |
| 12 | One | 29.07.23 | 15:00 | min:15.1<br>max:24.6<br>avg:19.2 |  |
| 13 | One | 30.07.23 | 15:00 | min:15.7<br>max:21.6<br>avg:18.3 |  |

**Table A.4:**
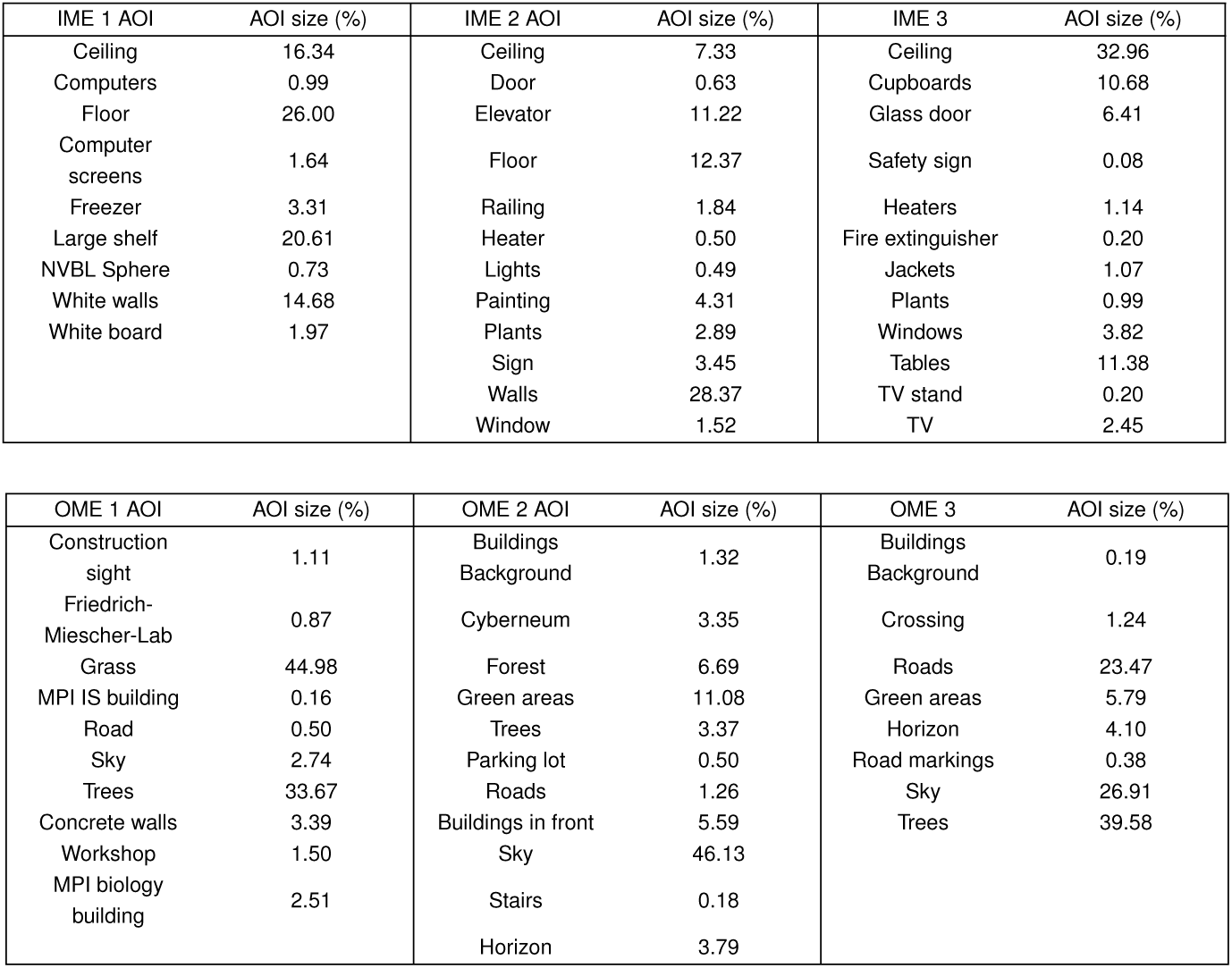
Sizes of AOIs (percentages of whole image size) in each measurement environ- ment. The AOIs were extracted from images of the scenes (Fig. B.7).

